## Supplementary figures for "Chromatin activity of IκBα mediates the exit from naïve pluripotency"

### Figure S1 Related to Figure 2

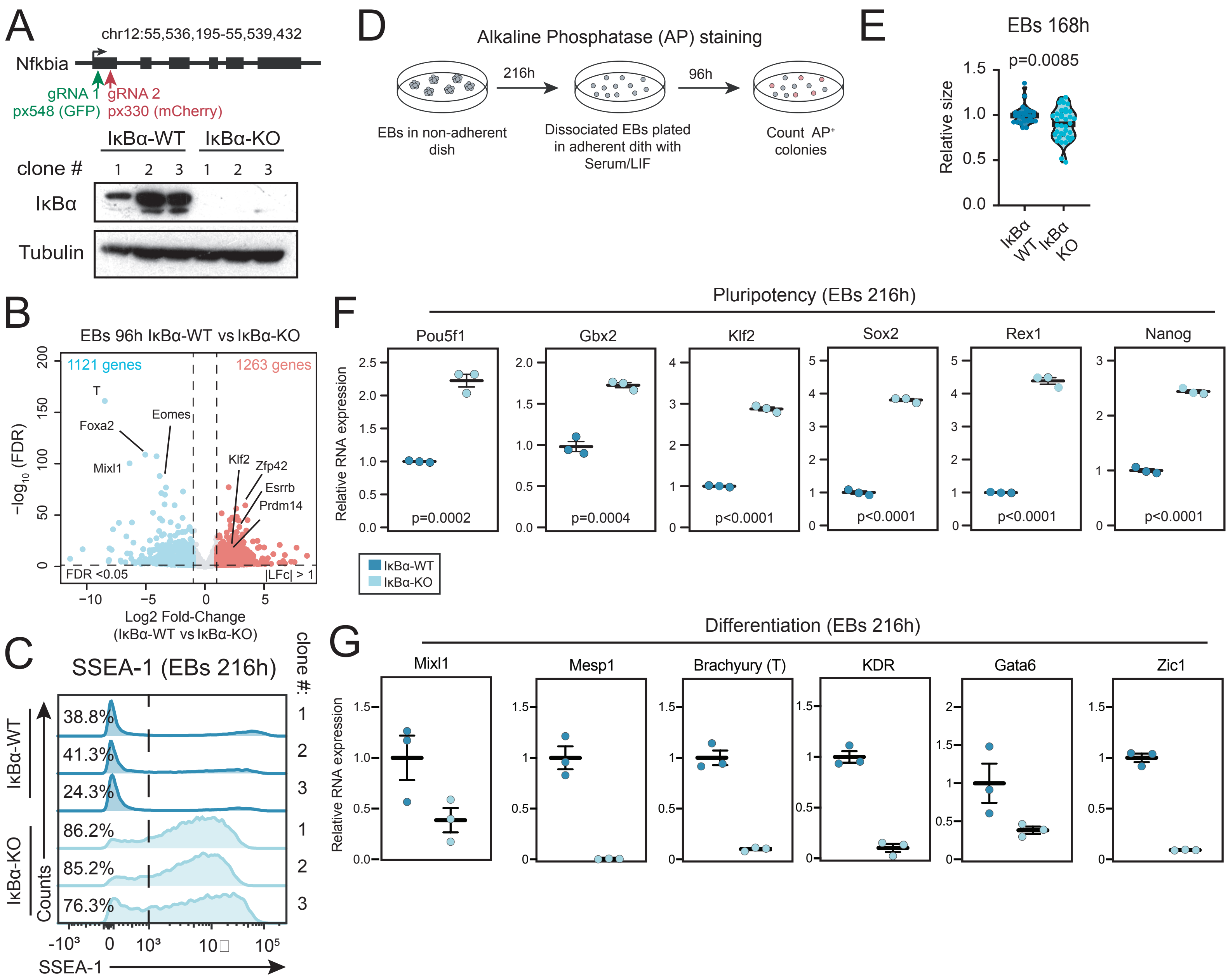

### Figure S2 related to figure 3

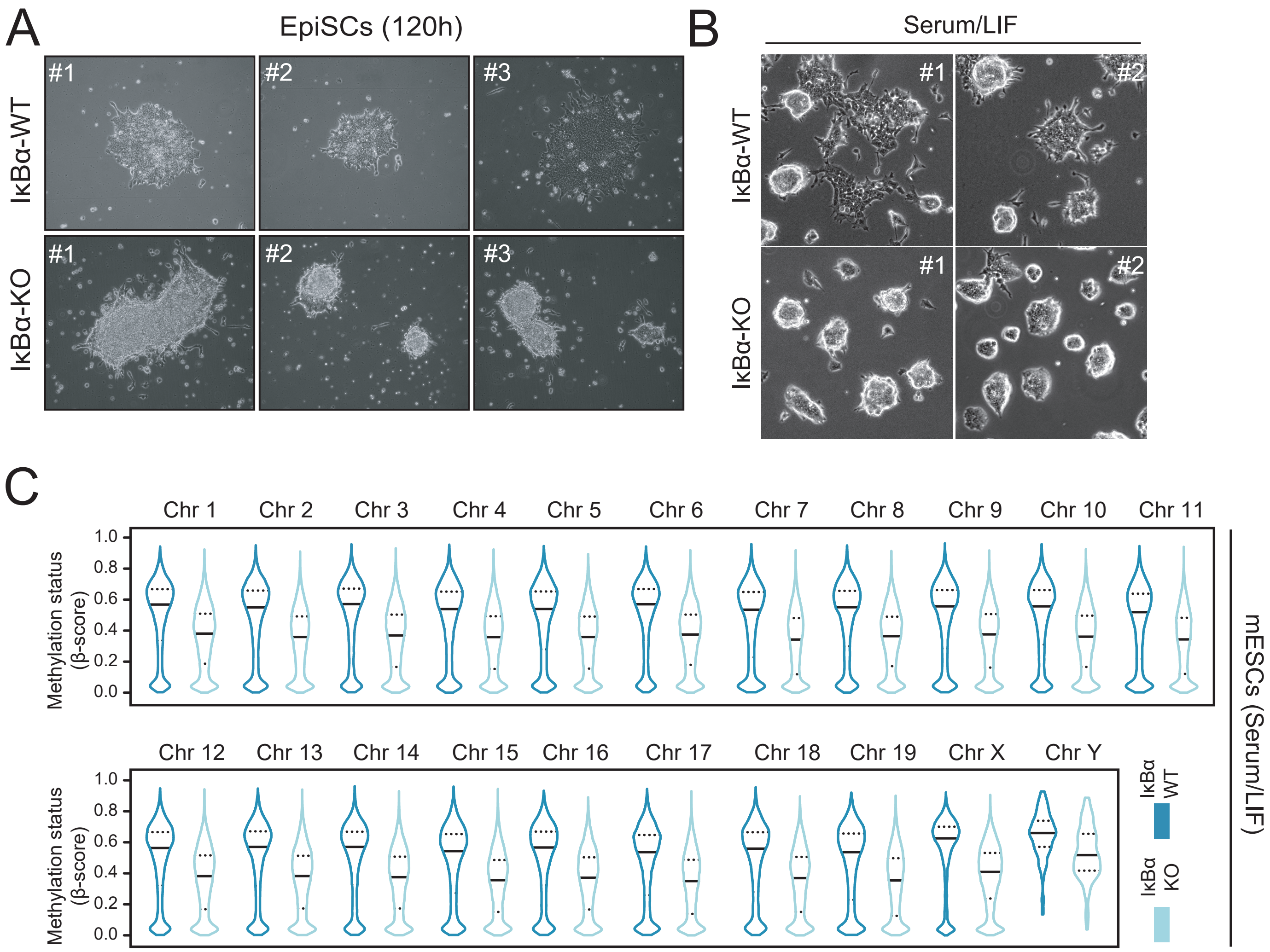

### Figure S3 related to figure 4

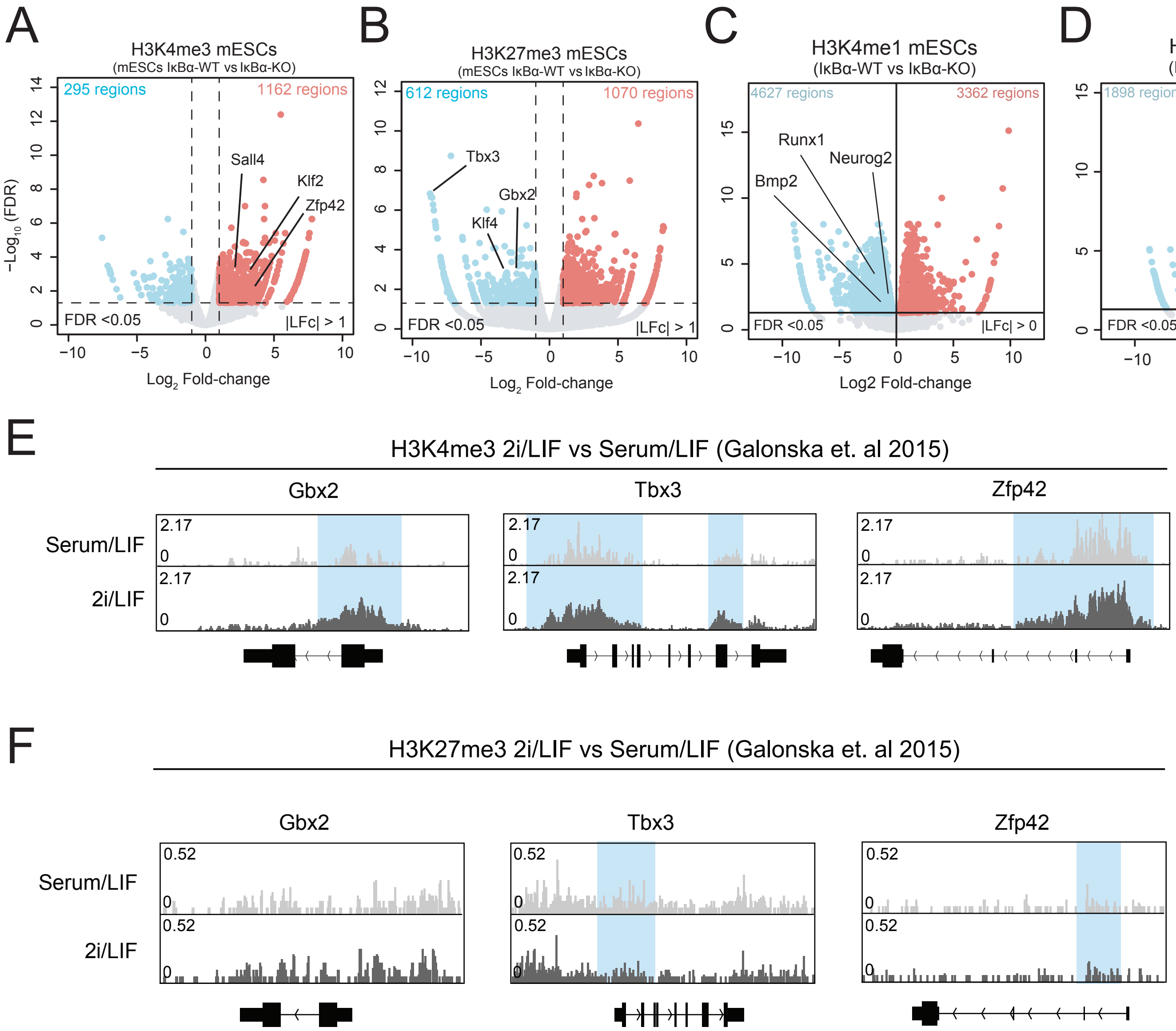

### Figure S4 related to figure 5

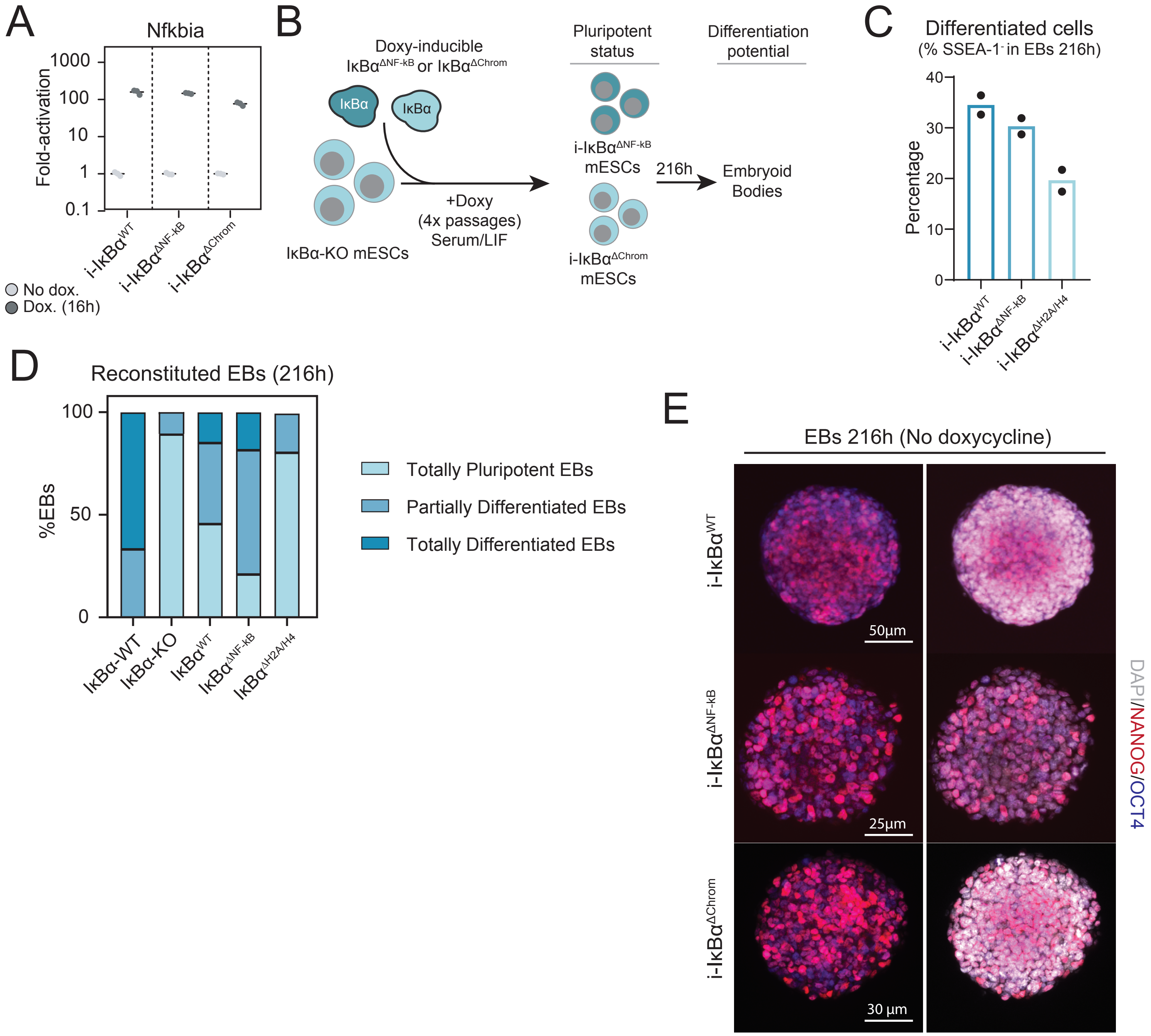
